# Introductory notes before zebra finch song have unique timing properties while sharing acoustic properties with song

**DOI:** 10.1101/2025.05.19.654952

**Authors:** Divya Rao, Raghav Rajan

## Abstract

Preparatory neural activity precedes the initiation of simple movements and a key feature of this preparatory activity is its trial-by-trial correlation with features of the upcoming movement. Recent studies in the zebra finch, a songbird with a complex, naturally learned, movement sequence (song), have suggested that the repeats of short introductory notes (INs) at the start of each song bout, reflect motor preparation. However, whether IN properties correlate with upcoming song features remains poorly understood. Here, we addressed this question by recording and analyzing male zebra finch songs over a 3 year period. We found bout-to-bout correlations in the acoustic features of the last IN and the first song syllable. However, similar correlations were present between the first song syllable and the first IN and the first and second song syllable suggesting that INs are also part of the song sequence. Next, we found an age-related increase in the mean IN number before song and song tempo. If INs reflected preparation of song parameters, we expected age-related song changes to be predicted by IN-song correlations at a younger age. We did not find any such correlations. Finally, we compared INs to other repeated syllables within song and outside song bouts and found that the speeding up of intervals between successive INs is unique to INs. Overall our results showing similarities in the acoustic features of INs and song syllables suggest shared neural control of INs and song syllables, while differences in timing suggest different neural mechanisms controlling IN timing.

**Significance Statement:** Simple movements are believed to be “prepared” in the brain before execution and this preparatory neural activity is correlated with features of the upcoming movement. Recent studies have suggested that the short introductory notes before the complex song sequence of the zebra finch reflect motor preparation. Whether correlations exist for introductory notes and upcoming song remains poorly understood. Here we found bout-to-bout correlations between the acoustic properties of introductory notes and the first song syllable but our analyses suggest that this reflects shared neural control of the acoustics of both introductory notes and song syllables. We also found differences in the timing of introductory notes suggesting different neural mechanisms for controlling introductory note timing.

## Introduction

How does the brain initiate a movement? Current research suggests a “preparatory” period before movement initiation when neural activity converges on a consistent initial state (Churchland et al., 2006b, 2010; Shenoy et al., 2011; Li et al., 2016; Svoboda and Li, 2018). Support for this hypothesis comes from delayed reaching tasks where subjects (humans or other animals) are trained to execute simple movements like reaching for an object. At the start of a trial, an instruction stimulus is provided that tells subjects where to move, but subjects are expected to withhold their movements until a “GO” cue is provided. The “GO” cue is presented after a variable delay period from presentation of the instruction stimulus.

The reaction time, measured as time between GO cue presentation and movement initiation, is shorter when the delay period is longer suggesting the need for a time to “prepare” the movement (Rosenbaum, 1980; Riehle and Requin, 1989; Churchland et al., 2006b). Premotor neural activity, recorded during this task, shows changes during the delay period (Churchland et al., 2006b; Guo et al., 2014) and an important feature of delay period neural activity is a reduction in variability across trials; more variable activity at the beginning of the delay period to less variable activity just before movement onset (Fig. 1A). Thus, for simple reaching movements, the brain appears to prepare by bringing premotor activity to a consistent initial state from which the correct patterns of movement related activity are produced immediately after the GO cue (Shenoy et al., 2011; Svoboda and Li, 2018).

**Fig. 1.**
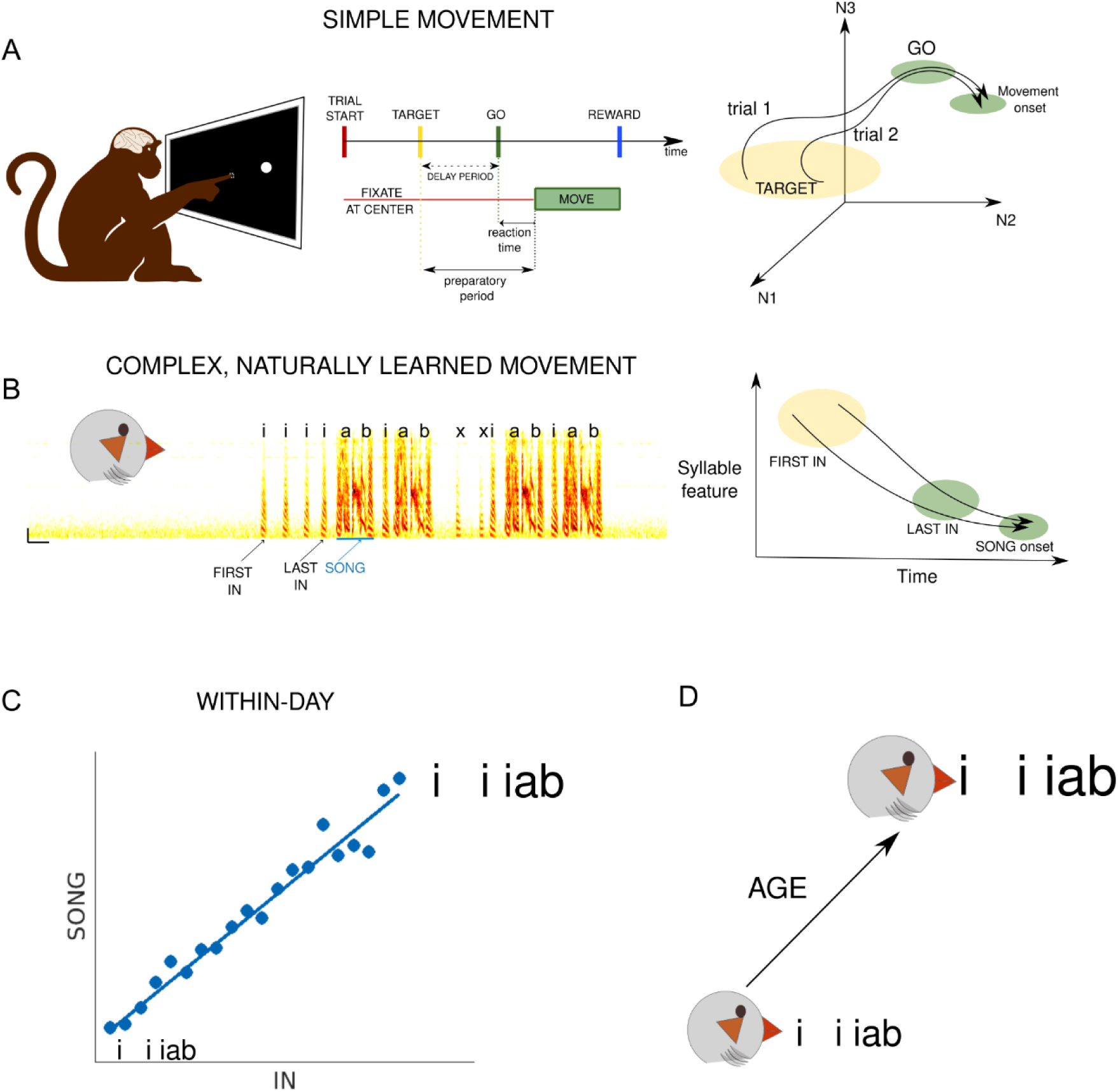
Preparatory activity and its relationship to upcoming movement. (A) Simple movement in monkeys: movement of finger from a central fixation point to a target as part of delayed reaching tasks. The neural activity on each trial (represented by the black traces), during the preparatory period, converges on a consistent point before movement onset. (B) Zebra finch song is an example of a complex, naturally learned movement. ‘i’s represent Introductory Notes (INs), ‘ab’ repesent song sequence. Properties of INs converge on a consistent state before the onset of first song syllable ‘a’. (C) If INs represent preparation for song, properties of INs should be correlated to properties of upcoming song. ‘iiiab’ represent IN-SONG sequence, and size of letters represent features that are correlated within a day. (D) If INs represent preparation for song, age-related changes to INs and SONGs should be predicted by within-day correlations.

What is the nature of motor preparation before more complex, naturally learned, motor sequences? A well-studied example of a complex, naturally learned, motor sequence is the song of the adult male zebra finch, a songbird (Fee and Scharff, 2010). Song is a sequence of sounds interleaved by silent gaps (Fig. 1B, ’a’ and ’b’ represent the song sequence). It is part of the courtship ritual of the male zebra finch (Sossinka and Böhner, 1980; Zann, 1996; Fee and Scharff, 2010) and is naturally learned by young birds (Immelmann, 1969; Price, 1979; Zann, 1996). Song bouts typically begin with a variable number of repetitions of a short sound called an introductory note (IN marked as ’i’ in Fig. 1B) (Price, 1979; Sossinka and Böhner, 1980). As INs repeat, the timing and acoustic features change; intervals between successive INs become shorter and more stereotyped and the acoustic features reach a stereotyped “state” just before the start of the first song (Rajan and Doupe, 2013). This progression is unaffected immediately after deafening or peripheral nerve cuts (Rao et al., 2019). This sensory-feedback independent progression from a variable first IN to a stereotyped last IN is similar to changes in delay-period neural activity suggesting that INs reflect motor preparation for song initiation.

An important feature of neural preparatory activity is its correlation with features of the upcoming movement. In primates, preparatory neural activity is different for different movements; for eg. preparatory neural activity occupies different initial states for fast and slow reaching movements. It also correlates with movement parameters on a trial-by-trial basis; within all fast reaching movements, preparatory neural activity is correlated with movement speed on that trial (Churchland et al., 2006a). Whether zebra finch INs also correlate with the upcoming song in a similar fashion remains poorly understood. Here, we addressed this by recording and analyzing songs from multiple adult, male, zebra finches across a 3 year period. Specifically, we first tested for the existence of bout-to-bout correlations between IN properties and song properties (Fig. 1C). Second, previous studies have shown age-related changes in song properties (Brainard and Doupe, 2001; Pytte et al., 2007; Glaze and Troyer, 2013; James and Sakata, 2019). If IN properties are correlated with song properties, one would also expect age-related changes in IN properties. Additionally, bout-to-bout correlations between INs and song properties, at a younger age, should predict age-related changes in IN and song properties (Fig. 1D). We tested for these by analyzing songs from the same birds at multiple different ages.

## Materials and Methods

All experiments at [Author University 1] were approved by Institute Animal Ethical Committee in accordance with the guidelines of the Committee for the Control and Supervision of Experiments on Animals [Author Country]. All song recording procedures for songs recorded at [Author University 2] were approved by the [Author University 2] Institutional Animal Care and Use Committee in accordance with NIH guidelines. We recorded songs from a total of 46 adult male birds (age > 90 days post hatch) with some purchased from an outside local vendor (n=19) and some bred at [Author University 1] (n=21) or [Author University 2] (n=6). The age of all purchased birds was assumed to be 60 days post hatch (dph) as they had red beaks (beaks are black for birds < 60 dph). At times when birds were not being recorded, they were housed in large cages with 5-8 other birds in a bird colony maintained with a 14h/10h light/dark cycle. Food and water were provided at all times.

### SONG RECORDING

Birds were isolated from the colony and placed in separate cages in a sound attenuation enclosure (NewTech Acoustic Systems, Bangalore) maintained at 14h light / 10 h dark cycle. A microphone (AKG Acoustics C417PP) was clipped on the roof of the cage to record song. Song recordings were either in ‘triggered’ or ‘continuous’ mode. Briefly, in ‘triggered’ mode periods of recordings that crossed a pre-set threshold were saved along with an additional 1-3 seconds of data flanking this period on either side. In ‘continuous’ mode, audio files were saved continuously for the entire recording period. All data was recorded and saved to disk at 44100 Hz sampling rate using custom written software (Python or Matlab). The different sets of birds used for different analyses and the overlap between these sets are explained below.

### BIRDS USED FOR ANALYSIS OF IN-SONG CORRELATIONS

20 birds were recorded on multiple days (median 3 days [range 2-8 days] sessions - put median sessions here done) in the age range from 89 - 1087 dph. As far as possible, we maintained the same position of the microphone for a given bird on different days of recording. A subset of birds and sessions (2 session each from 14/20 birds) overlapped with those analyzed for day-to-day changes in IN properties in a previous study. One bird (1/20) overlapped with the birds used for ts-cut surgery in that same study, but the recording sessions analyzed here are different and are from well before the surgical procedure. A subset of birds (5/20) were recorded earlier at [Author University 2] and have sessions from two nearby days, <5 days apart, included in this analysis. For characterizing IN changes towards song (Fig. 2) and bout-to-bout correlations of INs and song (Table 1 and Fig. 3) songs recorded from one session (<1 yr of age) in each bird were analyzed. Multiple sessions were analyzed for age related changes in INs and songs (Fig. 4-6). The difference between days of recording spanned a wide range (1-812 days) and the time of recording varied across sessions for the same bird.

**Fig. 2.**
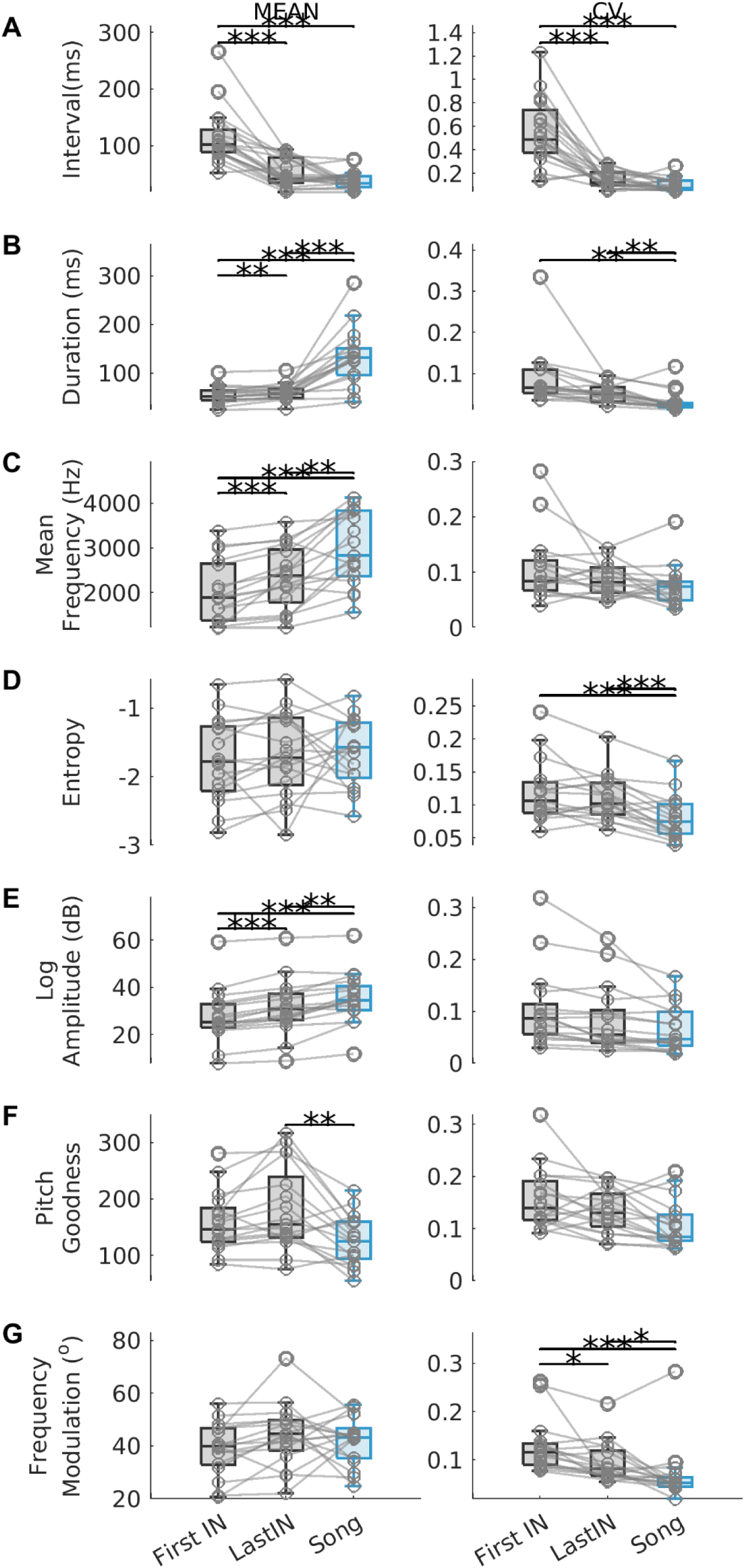
Introductory Note (IN) properties change in the direction of the song. (A-G) Comparison of properties between First and Last IN position (gray) and the first syllable of the upcoming song (blue). Circles joined by lines represent data from individual birds, boxes represent group data across birds. Column 1 shows mean and column 2 shows CV or Coefficent of Variation for intervals (A), syllable acoustic properties namely duration (B), mean frequency (C), entropy (D), log amplitude (E), pitch goodness (F), frequency modulation (G) respectively. * denotes p ≤ 0.05, ** p ≤ 0.01, *** p ≤ 0.005, Repeated Measures ANOVA followed by post-hoc Tukey-Kramer test.

**Fig. 3.**
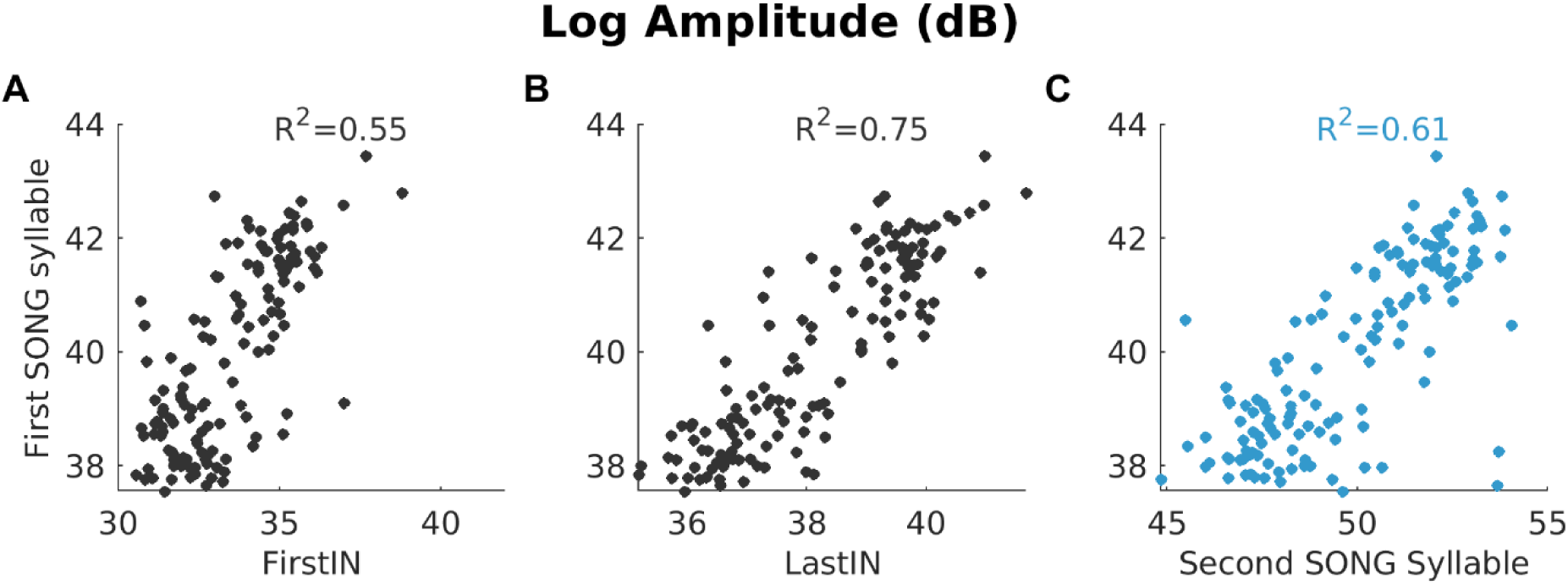
Trial-to-trial correlations between acoustic properties of IN and song. (A-C) Example session from one bird that shows significant positive correlations for Log Amplitude of first syllable in the (first) motif along y-axis with (A) First IN position, (B) Last IN position and (C) second syllable respectively along x-axis. Circles represent data from individual trials or bouts. *p* ≤ 0.05, Pearson correlation coefficient calculated after removing outliers.

**Fig. 4.**
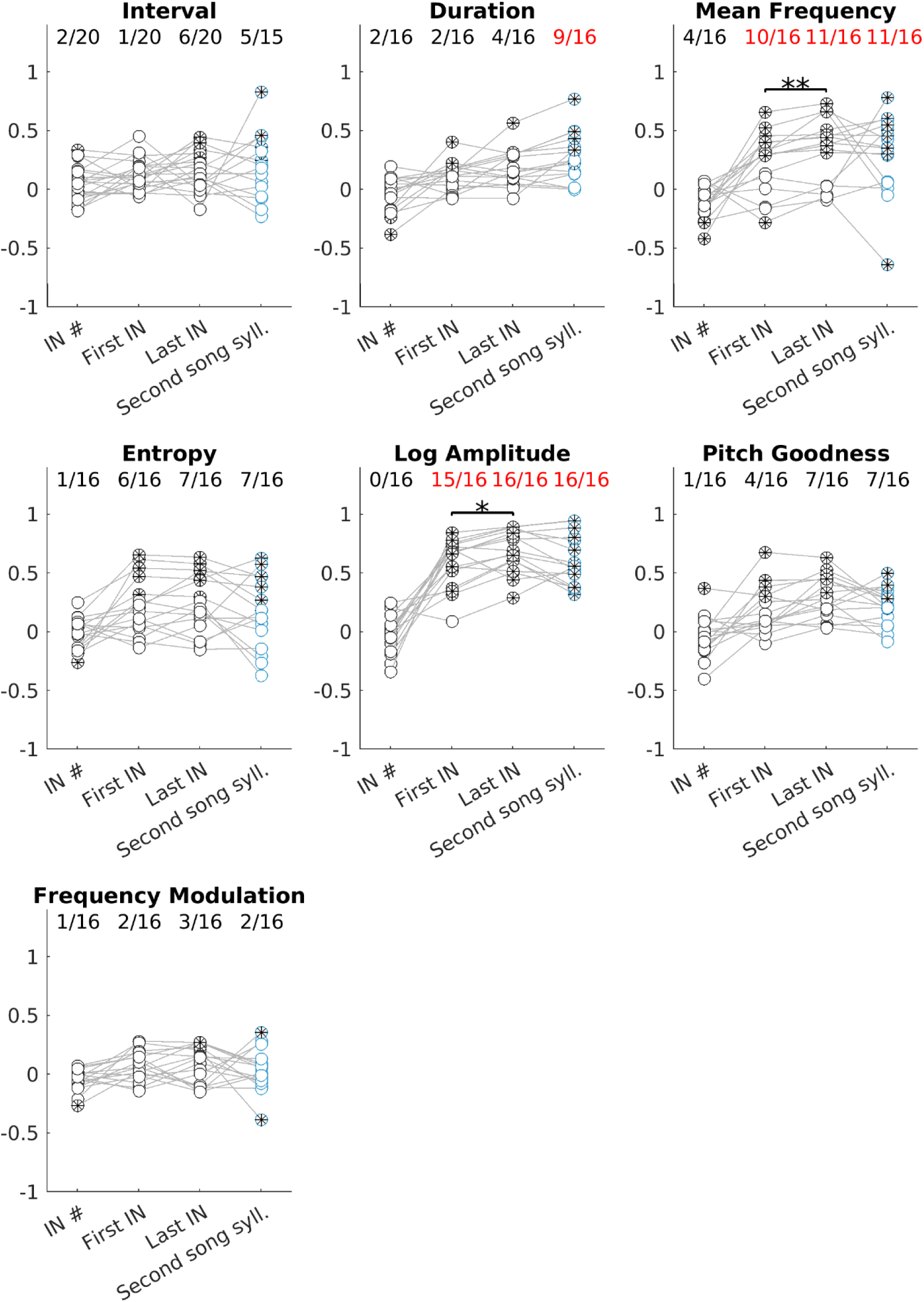
Correlations with first song syllable. IN number, timing and acoustic properties (duration, mean frequency, entropy, log amplitude, pitch goodness and frequency modulation for a syllable) at First IN, Last IN and second song syllable positions were correlated to timing and acoustic properties at first song syllable position across bouts. Each circle represents r-values for bout-to-bout correlations in one bird. Lines connecting circles represent data from the same bird bird. circles with a ’*’ represent significant correlations and unfilled circles represent non-significant correlations. Proportion of birds with significant correlations are indicated on top and marked with red if proportions were ≥ 0.5. A property is considered significantly correlated if proportions are ≥ 0.5. Correlation strengths across pairs of groups were compared using sign-rank by including only significant correlation values. Correlations to First IN were compared with correlations to last IN, correlations to last IN were compared with correlations to second song syllable. * denotes p ≤ 0.05, ** denotes p ≤ 0.01, Wilcoxon signed-rank test.

**Fig. 5.**
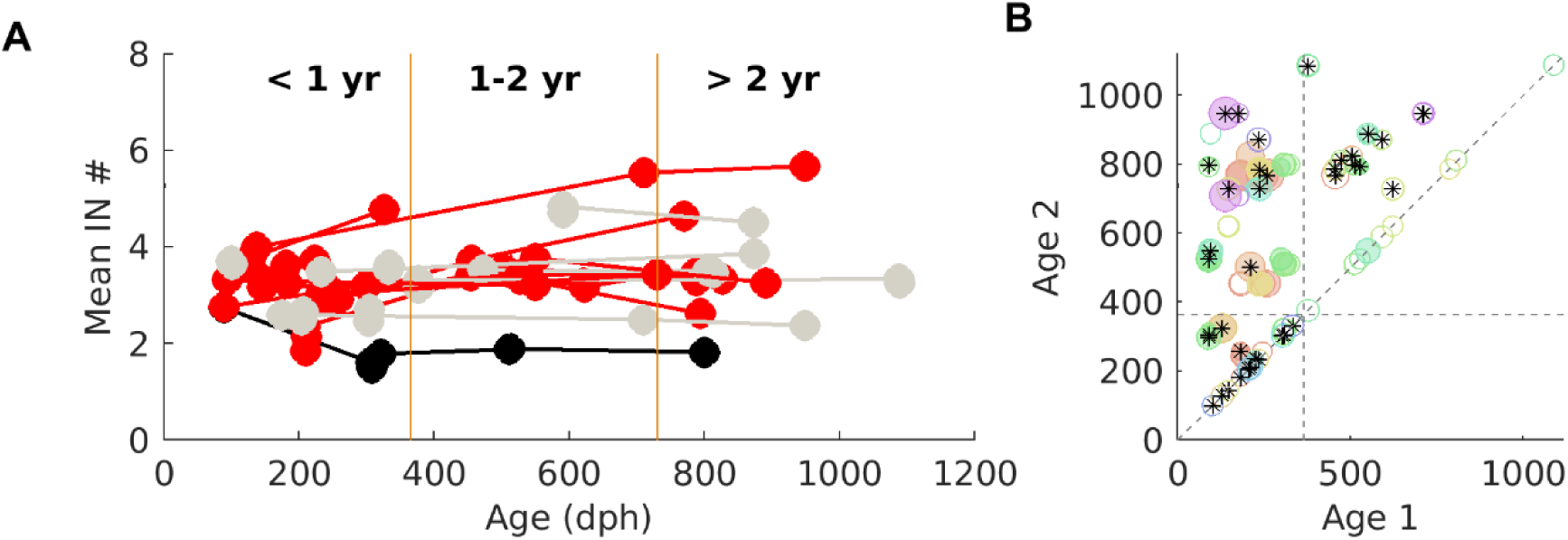
IN number increases in the first year of age. (A) Mean number of INs recorded at different ages across birds. Each circle represents the mean IN number for one session and lines join multiple datapoints from the same bird. Vertical yellow lines mark 1 year (365 dph) and 2 year (730 dph) respectively. Red lines indicate significant increase, black lines indicate significant decrease and gray lines indicate no change with age for a bird. (B) Circles represent comparison between an earlier day (Age 1 along x-axis) to a later day (Age 2 along y-axis) of recording. Horizontal and vertical gray dashed lines mark 1 year (365 dph). Diagonal gray dashed lines represent Age 1 = Age 2. Size of the circles represent change in mean IN number between Age 1 and Age 2 and filled circles represent a significant change. Different colors represent different birds. Points marked with star (*) were selected for maximum change in IN number for a bird in an age group that are also plotted in (6C). KruskalWallis ANOVA in (A) for individual birds followed by Tukey-Kamer test to mark significant pairs of days in (B), *p* ≤ 0.05 considered significant.

**Fig. 6.**
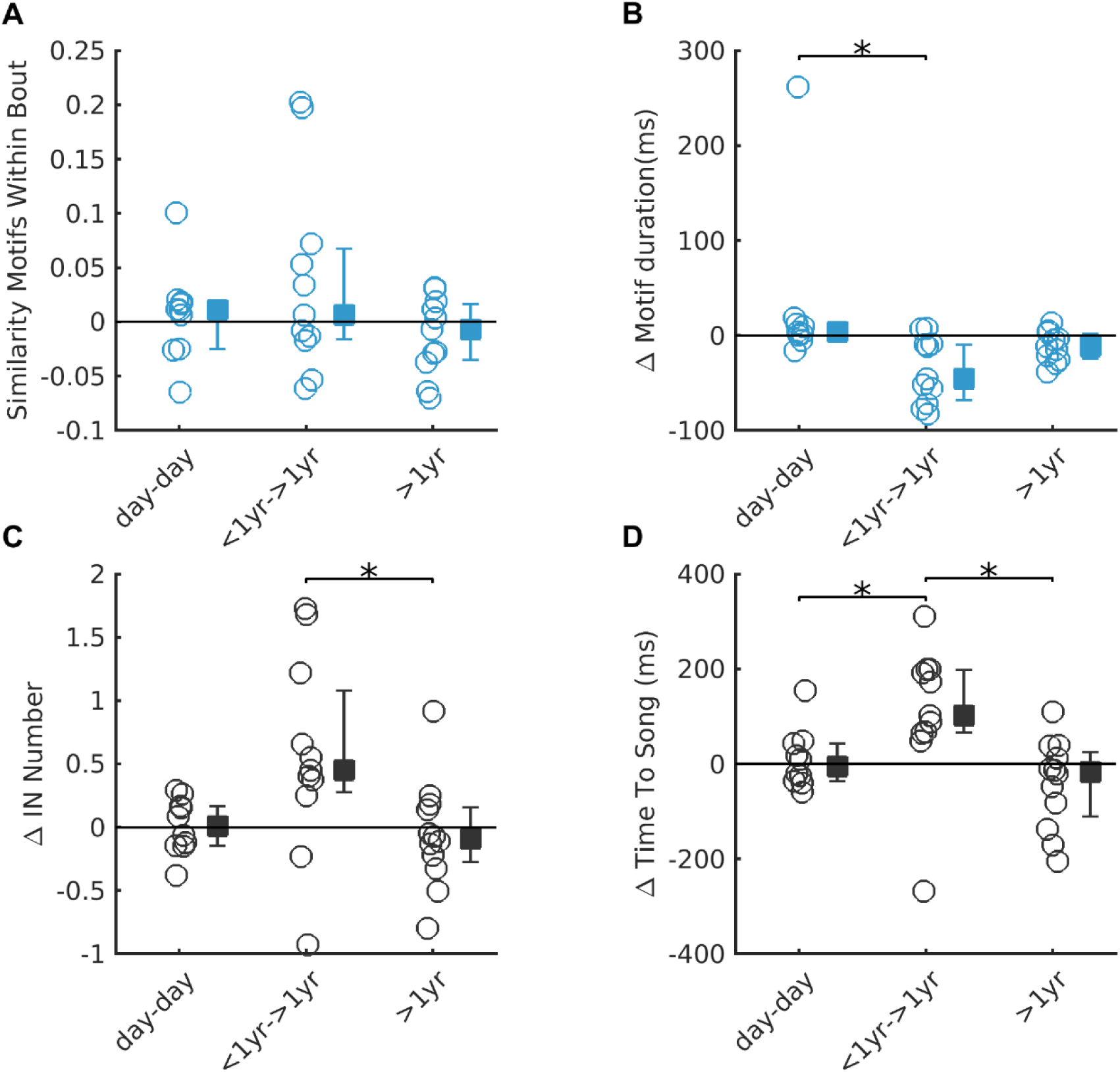
Song and IN properties simultaneously change in the first year of age. (A - D) Changes in Motif similarity (A), motif duration (B), IN number (C) and time-to-song (D) are plotted across different age categories. Circles represent change in mean property between pairs of days for individual birds. All pairs of days are the same pairs of days selected for maximum change in IN number in Fig. 5. Square and whiskers represent median and interquartile range for the 3 age groups. * denotes *p* ≤ 0.05 KruskalWallis ANOVA followed by Tukey-Kramer test.

**TABLE 1.** Correlation between IN and song syllables. Timing and acoustic properties of first song syllable (see Row Names) were correlated to IN number, or corresponding timing and acoustic properties for first IN position, last IN position or second song syllable (see Column names). Pearson correlation coefficient was calculated after removing outliers for one session in each bird and considered significant if p ≤ 0.05. Each cell indicates the proportion of birds with significant correlations along with the range of R-squared values for significant correlations, within parenthesis. Highlighted cells indicate properties for which significant correlation coefficients were observed in at least half of the birds. The subsequent columns repeat the analysis for birds with head-implanted microphone and ts-nerve cut birds.

|  |  | Unmanipulated birds |  |  |  |  |  |  |  | Surgically manipulated birds |  |  |  |
| --- | --- | --- | --- | --- | --- | --- | --- | --- | --- | --- | --- | --- | --- |
|  |  | Suspended microphone (n=20) |  |  |  | Head fixed microphone (n=4) |  |  |  | Ts Cut (n=8) |  |  |  |
|  |  | IN number | First IN | Last IN | Second Song syllable | IN Number | First IN | Last IN | Second Song syllable | IN Number | First IN | Last IN | Second Song syllable |
| First Song Syllable | Interval | 2/20 [0.027564 - 0.11579] | 1/20 [- 0.040692 -] | 6/20 [0.055634 - 0.20126] | 5/15 [0.062064 - 0.69446] | 0/4 [-] | 1/4 [- 0.094595 -] | 0/4 [-] | 0/3 [-] | 2/7 [0.020996 - 0.084361] | 1/7 [- 0.060756 -] | 3/7 [0.077037 - 0.20297] | 5/6 [0.059212 - 0.31928] |
|  | Duration | 2/16 [ 0.055525 - 0.14306] | 2/16 [ 0.050712 - 0.16412] | 4/16 [ 0.07711 - 0.32075] | 9/16 [ 0.049804 - 0.59583] | 0/4 [-] | 1/4 [- 0.10509 -] | 0/4 [-] | 1/4 [- 0.41001 -] | 2/8 [0.069903 - 0.094664] | 5/7 [ 0.019376 - 0.32247] | 3/7 [ 0.11523 - 0.15811] | 5/8 [ 0.066957 - 0.32461] |
|  | Mean Frequency | 4/16 [ 0.037527 - 0.17197] | 10/16 [ 0.07751 - 0.43468] | 11/16 [ 0.097598 - 0.53551] | 11/16 [ 0.086829 - 0.61189] | 0/4 [-] | 0/4 [-] | 2/4 [ 0.2547 - 0.26094] | 3/4 [ 0.18096 - 0.2585] | 2/8 [0.026057 - 0.035218] | 6/7 [ 0.085925 - 0.17935] | 6/7 [ 0.085449 - 0.32444] | 7/8 [ 0.17162 - 0.59137] |
|  | Entropy | 1/16 [- 0.065818 -] | 6/16 [ 0.043557 - 0.43134] | 7/16 [ 0.089148 - 0.40611] | 7/16 [ 0.033528 - 0.39943] | 0/4 [-] | 0/4 [-] | 2/4 [ 0.21301 - 0.23058] | 2/4 [ 0.13476 - 0.37133] | 3/8 [0.033457 - 0.098931] | 4/7 [ 0.075541 - 0.35578] | 5/7 [ 0.074278 - 0.53467] | 7/8 [ 0.077904 - 0.69269] |
|  | Log Amplitude | 0/16 [-] | 15/16 [ 0.1022 - 0.71674] | 16/16 [ 0.086365 - 0.80228] | 16/16 [ 0.10257 - 0.90177] | 0/4 [-] | 1/4 [- 0.25684 -] | 3/4 [ 0.095298 - 0.29403] | 3/4 [ 0.10264 - 0.23485] | 3/8 [0.070799 - 0.091607] | 7/7 [ 0.18765 - 0.64247] | 7/7 [ 0.43535 - 0.83514] | 8/8 [ 0.46317 - 0.78988] |
|  | Pitch Goodness | 1/16 [- 0.13814 -] | 4/16 [ 0.093954 - 0.45981] | 7/16 [ 0.082209 - 0.40086] | 7/16 [ 0.039365 - 0.25201] | 0/4 [-] | 1/4 [- 0.55047 -] | 0/4 [-] | 0/4 [-] | 3/8 [0.022745 - 0.073854] | 6/7 [ 0.1454 - 0.69199] | 7/7 [ 0.16424 - 0.82966] | 8/8 [ 0.31519 - 0.88773] |
|  | Frequency Modulation | 1/16 [- 0.070169 -] | 2/16 [ 0.039416 - 0.079888] | 3/16 [ 0.054115 - 0.075381] | 2/16 [ 0.12817 - 0.14779] | 0/4 [-] | 0/4 [-] | 0/4 [-] | 1/4 [- 0.14862 -] | 2/8 [0.053764 - 0.067706] | 4/7 [ 0.042636 - 0.16752] | 3/7 [ 0.099547 - 0.36539] | 5/8 [ 0.16071 - 0.22946] |

### DATA OVERLAP WITH PREVIOUS STUDIES

Songs of birds from [Author University 2] were recorded as part of a different study characterizing INs. Besides characterizing the properties of INs in every sequence and the associated neural activity during INs, the study also correlated the similarity of pairs of last INs to pairs of first song syllables. However, the direct relationship between individual properties of INs and songs was not compared. Here, we used these songs to directly compare the properties of INs and songs.

The IN-song correlation analysis was repeated for one session of undirected song each from 5 birds recorded earlier with head-implanted microphones for a different study. Head-implanted microphone recordings control for amplitude measurement differences in suspended microphone recordings, that may arise due to changes in relative position of the bird and the microphone across trials. IN-song correlations of acoustic properties with head-fixed microphones were analyzed as a control to rule out the possibility that IN-song correlations were a result of differences in relative position of the bird from the microphone. IN-song correlations have not been analyzed as part of the original study.

IN-song correlation analysis was repeated for one session of undirected song each from 8 birds recorded after ts-nerve (tracheosyringeal nerve) cut surgery as part of an earlier study. This data provided an advantage to look at IN-song acoustic correlations in the absence of properties related to syllable identity as these are lost following ts-nerve cut manipulation. IN-song correlations were not analyzed as part of the original study.

### DATA FOR REPEAT SYLLABLES

Songs of 17 birds in the colony were identified that specifically repeated at least one syllable within the motif. One session each from these birds was used to compare the syllable repetition properties among the different types of repeated syllables produced by the bird namely INs, motif syllable repeats and calls. One session from one bird was also analyzed for IN-song correlations. One of the birds overlapped with ts-nerve cut birds, but the songs analyzed were from before surgery. One of the birds recorded in this set was also bred and recorded earlier at [Author University 2].

### DATA ANALYSIS

All the analyses were performed using custom-written scripts in MATLAB (www.mathworks.com). Audio files were processed and vocalizations were labeled. Briefly, audio files were segmented into syllables based on a user-defined amplitude threshold, syllables were defined as segments greater than 10 ms and inter-syllable gap shorter than 5ms were merged. Labels were assigned to syllables in a semi-automatic manner. Automatic labels were assigned using a modified template-matching procedure (Glaze and Troyer, 2006) or Klustakwik clustering (http://klustakwik.sourceforge.net/) of acoustic features calculated using Sound Analysis Pro (https://soundanalysispro.com/matlab-sat). Labels were then manually checked for all files. Files were split into bouts and all bouts containing songs (motifs) were selected for analysis. Typically these bouts consisted of repeating IN sequences before the first song.

#### Defining a bout and bout interval criteria

A bout was defined as a period of vocalizations separated by 2 seconds of silence. This criteria was applied to select bouts from recordings in ‘continuous’ mode. However, many audio files recorded in ‘triggered’ mode did not have 2 seconds of silence before the first syllable in the file. This occurred when initial vocalizations in the bout were soft and the set trigger threshold was crossed by a later vocalization. Such audio files had less than 2 seconds of silence before the first syllable in the file. However, we assumed that there was silence before the start of the file as the set trigger-threshold was not crossed and so we included such files in our analysis. The bout criteria for triggered recordings was reduced to include enough audio files (>15 bouts) for analysis and the criteria ranged from 500-1500 ms only for the beginning of the file. Within a file, we still considered 2000ms of silence and we always considered 2000ms of silence for continuous recordings. across birds. For a given bird, we maintained the same bout criteria across sessions. This was important as we observed that mean IN number depended on bout interval criteria; as shown previously, mean IN number reduced a little when considering a bout criteria of < 2s (Sossinka and Böhner, 1980; Rajan and Doupe, 2013). However, mean IN number between days was not different when the same bout interval criteria was applied.

#### Determining minimum number of bouts for IN analysis

The number of INs varied across bouts and the number of bouts sung by birds varied across sessions and birds (median 73.5 song bouts: range 4 - 430). It was important to characterize how the number of bouts affected the estimate of mean IN number for a session, and to determine the minimum number of bouts beyond which the estimate did not change considerably. For all sessions with >100 song bouts (30 sessions), a range of different number of bouts were sub-sampled and the corresponding average standard deviation across 1000 iterations of Monte Carlo simulations was calculated. The minimum number of bouts for IN analysis was identified as the number of bouts beyond which increasing the number of bouts did not change the average standard deviation by more than 0.01. The standard deviation met our criteria by 10 bouts for 18/30 sessions and by 15 bouts for 28/30 sessions. Hence, across all days and birds, only sessions with at least 15 song bouts were selected for IN analysis. This excluded two sessions from a bird (1/20) with < 15 song bouts.

#### Calculating IN and song properties

For analyzing IN and song properties across trials, only INs in the beginning of the bout and the first motif, that followed immediately after the INs, was considered, unless otherwise specified. For each song bout, the number of INs was calculated preceding the first motif. The timing and acoustic properties of INs and song were calculated as follows: timing of IN for first position or first interval was the time interval between offset of first IN and onset of second IN. Timing of IN for last position or the last interval was calculated as the time interval between offset of last interval and the onset of the first song syllable. Song timing for the first position in the motif was calculated as the interval between offset of first motif syllable and onset of second motif syllable. Song timing for the second position in the motif was calculated as the interval between offset of second motif syllable and onset of third motif syllable. Similar to timing, the acoustic properties of INs were calculated for first and last IN and for the first and second motif syllable. For comparisons of acoustic properties involving changes in IN from first to last position, birds with two types of INs in the IN sequence were excluded as the syllable identity in these birds would be different for first and last IN. For comparisons of intervals involving first and second song position, birds with only two syllables in the motif were excluded as there were no second song intervals.

#### Acoustic properties of syllables

The acoustic properties for IN and song syllables were measured as described by Sound Analysis Pro 2011 (Tchernichovski et al., 2000) (http://soundanalysispro.com/) using SAP matlab code (https://soundanalysispro.com/matlab-sat). These basic features reduce dimensionality of the complex sound spectrogram for analysis and make it easier to understand the changes in different aspects of sound. A brief intuitive understanding of the features is described below:

- Log Amplitude: It measures the loudness of a syllable. The intensity or power in audio signal is measured relative to an arbitrary baseline for silence. It is reported in log scale units or dB.
- Duration: Time from onset to offset of the syllable. Onset and offset are determined based on an amplitude threshold that distinguishes sound from silence in the audio signal. It is measured in units of seconds or milliseconds.
- Mean Frequency: Audio signal can be decomposed into different frequencies present in the signal. The mean frequency is a pitch measure that assesses the center of the distribution of intensity across different frequencies. It is measured in Hz. Since mean frequency is dependent on intensity, it is an amplitude related property.
- Pitch goodness: It measures the periodicity of harmonic pitch (frequency stacks) in the syllable. Syllables with harmonics or ‘frequency stacks’ (as observed in a spectrogram) have higher value, and syllables that are noisy or pure tone have lower value.
- Weiner entropy or entropy: It measures the noisiness in the syllable. A noisy syllable appears as broadband or white noise in the spectrogram with intensity spread equally across all frequencies. White noise has a value of 1 and pure tone (sound intensity concentrated at one frequency) has a value of 0. However, this is converted to log-scale of 0 to minus infinity. More noisy syllables will have entropy 0. Since entropy is dependent on intensity, it is an amplitude related property.
- Frequency Modulation: It is an estimate of slope of the frequency trace on the spectrogram, measured in degrees. Steeper the slope, higher the modulation.

#### Analyzing age-related changes in IN and song properties

Changes in IN and song properties between two different days were measured by subtracting the mean or variability calculated for the earlier of the two days from those measured on the later of the two days. For changes in INs between days, in addition to properties related to IN number, IN timing and IN acoustic properties, the time-to-song was also calculated. Time-to-song was defined as the duration of the IN sequence from onset of the first IN to onset of the first song syllable. To quantify song changes with age, parameters of song that have earlier been reported to change with age were calculated. These included motif duration and song similarity (Brainard and Doupe, 2001; Pytte et al., 2007; Glaze and Troyer, 2013; James and Sakata, 2019). Motif duration was measured as the time from the onset of the first motif syllable to the offset of the last motif syllable. Song similarity was calculated as average similarity index between pairs of motifs from a recording day. Ten motifs of a fixed syllable sequence were randomly selected across all motifs, and each of the ten motifs was compared to the 9 other motifs using a similarity index algorithm (Mandelblat-Cerf and Fee, 2014), followed by taking the mean for all pairs. The similarity index gives a measure of how similar songs on a given day were to each other. Similar to other properties, changes between days was measured by subtracting the similarity index of an earlier day from a later day.

Other song properties related to song sequence and individual timing and acoustic features of first song syllable were also measured for change between days. To compare song spectral structure changes between pairs of days we measured the average similarity index of 10 randomly selected motifs on the two days. To compare song temporal structure between days the temporal pattern of the motif sequence was obtained by taking the amplitude profile and replacing this with ‘1s’ for the duration of syllables and ‘0s’ for the duration of intervals between syllables. Two patterns were compared up to the duration of the shorter sequence using cross correlation. Similar to comparison of spectral structure, 10 randomly selected motifs each from the two days were compared all-to-all and averaged. Sequence Consistency and Entropy were measured as described in an earlier study (James and Sakata, 2019). The analysis was restricted to the first motif sequence in the bout. For each motif sequence one additional syllable at both ends was included for calculating sequence consistency and entropy. Sequence Consistency (SC) measured the consistent transitions between syllables of the motif and was calculated as the ratio of most typical transitions divided by total transitions. Sequence Entropy(SE) measured the variability of syllable transitions, and was calculated as the sum of transition probabilities (p_i_) of all transitions from a syllable using the formula Σ-p _i_×log_2_(p_i_). Higher the value, more variable the transition. The change between pairs of days was measured by subtracting the values of an earlier day from a later day.

Age-related changes were calculated as changes in mean or variability of properties between pairs of days belonging to either of the 3 age-groups: (1) pairs of days < 5 days apart as short-term day-to-day change, (2) pairs of days with first day <1 year and second day > 1yr as long-term change in the first year and (3) pairs of days with both days > 1yr and more than 5 days apart as long-term changes after one year of age. If a bird was recorded for more than one day within an age-group, we compared all pairs of days with each other and for each age-group category, we chose pairs of days with the maximum change in IN number. We reasoned that this would give us the maximum chance of detecting changes, if there were any. The same pairs of days were used to calculate age-related changes in other IN and song properties to understand the extent of age-related changes in other properties when IN number changes were maximum.

#### Comparison of IN properties with motif duration and similarity within a session

To compare IN properties with motif duration and motif similarity within a day, we divided the bouts based on IN number at the start and then calculated the average motif duration and song similarity for bouts with same number of INs. To account for individual variation across birds, the number of INs was normalized to median IN number and motif duration was normalized to the corresponding median motif duration. We then asked whether song properties were different for the different IN numbers by comparing song properties across the different IN numbers. As another measure of number of INs, we also compared song changes with changes in the time-to-song, measured as the time from the start of the first IN to the start of the first motif syllable. For this, we split bouts into two groups based on the median value of time-to-song. The two groups were then compared across birds for differences in mean song properties corresponding to shorter or longer timing of INs. Other song properties chosen based on significant changes with age such as song sequence entropy, mean pitch goodness of first song syllable, and variability of frequency modulation of first song syllable were also compared corresponding to shorter or longer IN song timing. Similar analyses were done for comparing song properties corresponding to trials split based on properties of first or last IN position before song, namely lower and higher IN frequency modulation, less and more variable IN duration, less and more variable pitch goodness.

#### Analysis of repeat syllables

Repeat syllables belonging to either motifs, INs or calls were analyzed for each bird. Call repeats outside of song were considered as these were more common in all the birds we analysed. Similar to criteria used for INs in other datasets, repeat syllables with more than 15 instances were analysed. The number, interval and acoustic properties were analyzed using a similar procedure as that described for INs above. The ratio of intervals was defined as the current interval between successive repeats in the sequence divided by the next interval.

Shortening of intervals is denoted by a value of <1. Median ratio of intervals was measured across trials to represent data for a syllable. The acoustic distance to last repeat was defined as an inverse measure of similarity between a repeat syllable and all last repeat syllables. The six acoustic properties defined earlier were used to calculate acoustic distance, namely duration, mean frequency, log amplitude, entropy, pitch goodness and frequency modulation. To calculate acoustic distance, all repeat instances were randomly split in half. The last repeats from one half were chosen as the reference distribution occupying a six-dimensional space formed by the six acoustic properties. The Mahalanobis distance from this distribution to each repeat syllable in the second half was calculated to give the acoustic distance for each of these repeat syllables. Similar to ratio of intervals, ratio of acoustic distance was then calculated as the distance of every repeat divided by distance of following repeat syllable.

The median ratio of acoustic distance was measured across trials to represent data for a syllable. A median ratio of <1 denoted convergence towards the last repeat syllable.

### EXPERIMENTAL DESIGN AND STATISTICAL ANALYSIS

The experimental design of all groups tested for differences had comparable sample numbers. All comparison of independent groups of unequal sample sizes were tested for differences using non-parametric Kruskal-Wallis test. If the p-value was ≤ 0.05, Tukey-Kramer’s post-hoc test was used to identify pairs of groups that were significantly different These group comparisons included age-related changes in mean IN or song properties, age-related changes in IN number in individual birds, comparison of average song motif duration or song similarity for different IN numbers, and comparison of mean properties of repeat syllables belonging to either motifs, INs or calls. Differences between (more than two) bird-matched groups were tested using Repeated-Measures one-way ANOVA. If the p-value was ≤ 0.05, Tukey-Kramer’s post-hoc test was used to identify pairs of groups that were significantly different. This included comparison of mean or CV of syllable properties across bouts based on position, i.e. first IN, last IN and first song syllable. Paired-group comparisons were tested for differences using Wilcoxon signed-rank test with a p-value criteria of 0.05 for significant differences. These comparisons included significant correlation coefficient estimates of first IN - first song syllable with that of last IN – first song syllable, motif duration or similarity comparisons between shorter and longer time-to-song bouts, and properties of repeat syllables at first and last IN positions. All bout-to-bout correlations were measured using Pearson’s Correlation Coefficient of corresponding IN and song property values across bouts. Prior to measuring correlations, the outliers were removed (percentage removed - median- 2.19% data points; range - 0-28.99%). Outliers were detected as values beyond three times the median absolute deviation from the median for continuous variables and values beyond three standard deviations away from the mean for discrete variables. The correlations were considered significant if p-values were ≤ 0.05. The range of significant correlations were reported across birds along with proportion of birds significantly correlated.

### DATA AND CODE ACCESSIBILITY

All data and scripts for analysis are available on request from the corresponding author.

## Results

### Introductory note features change in the direction of upcoming song

Introductory notes (INs) are characterized by three properties; the number of INs before song at the start of each bout, their timing measured by the intervals between successive INs and their acoustic properties. As shown earlier (Rajan and Doupe, 2013; Rao et al., 2019), both the timing and the acoustic properties of INs changed systematically from the first to the last IN (Fig. 2). Additionally, changes in IN properties were in the direction of changes from INs to song. As INs progressed from the first IN to the last IN to the first song syllable, inter-syllable intervals got shorter (Fig. 2A, left, p < 0.001, Repeated Measures ANOVA, p < 0.001 for first IN vs song and first IN vs. last IN, post-hoc Tukey-Kramer test) and syllables progressively became longer (Fig. 2B, left, p < 0.001, Repeated Measures ANOVA, p < 0.001 for first IN vs. song and last IN vs. song, p = 0.02 for first IN vs. last IN, post-hoc Tukey-Kramer test), higher in frequency (Fig. 2C, left, p = 0.008, Repeated Measures ANOVA, p < 0.001 for first IN vs. song and first IN vs. last IN and p = 0.003 for last IN vs. song, post-hoc Tukey-Kramer test) and louder (Fig. 2E, left, p = 0.003, Repeated Measures ANOVA, p < 0.001 for first IN vs. song and first IN vs. last IN and p = 0.002 for last IN vs. song, post-hoc Tukey-Kramer test). In addition to changes in the mean, we also found a reduction in variability as INs approached song. Specifically, from the first IN to the last IN to song, the inter-syllable intervals became less variable (Fig. 2A, right, p = 0.048, Repeated Measures ANOVA, p < 0.001 for first IN vs. song and first IN vs. last IN, post-hoc Tukey-Kramer test), syllable duration (Fig. 2B, right, p =0.03, Repeated Measures ANOVA, p < 0.001 for last IN vs. song, p = 0.005 for first IN vs. song, post-hoc Tukey-Kramer test), entropy (Fig. 2D, right, p < 0.001, Repeated Measures ANOVA, p < 0.001 for first IN vs. song and for last IN vs. song, post-hoc Tukey-Kramer test) and frequency modulation (Fig. 2G, right, p = 0.002, Repeated Measures ANOVA, p < 0.001 for first IN vs. song, p = 0.04 for first IN vs. last IN and last IN vs. song, post-hoc Tukey-Kramer test) became less variable. Syllable duration (Fig. 2B, right) and entropy variability (Fig. 2D, right) were not significantly different from first IN to last IN (p > 0.05, Repeated Measures ANOVA). Overall, these results showed that the mean features of INs changed and became more similar to those of the upcoming song, as INs progress from the first to the last IN. In addition, timing variability systematically reduced from first to last IN, while the variability of acoustic features only reduced with the onset of song.

### IN number is not correlated with first song syllable features

If INs represent motor preparation for upcoming song, IN properties should be correlated with properties of the first song syllable on a bout-to-bout basis. We first correlated the number of INs with the properties of the first song syllable. In most of the birds, the number of INs (or the “time to song”, measured as the time between the first IN and the first song syllable) was not significantly correlated with either the interval between the first two song syllables or the acoustic properties of the first song syllable (Fig. 4, Table 1, p > 0.05, Pearson’s correlation co-efficient). Previous studies have shown that the number of INs (and the “time to song”) is positively correlated with the length of the interval between the first two INs and the acoustic similarity of the first IN to the last IN (Rajan and Doupe, 2013; Rao et al., 2019). Taken together with our results showing the absence of correlations between IN number and song features, this suggests that birds sing different number of INs to reach the same last IN state from different initial conditions (different first IN states).

### IN timing is not correlated with song syllable timing

We next examined correlations between the timing of INs and the timing of song. Specifically, we separately calculated correlations for first IN timing with song timing and last IN timing with song timing. If INs represent motor preparation, we expected an improvement in the IN-song correlations as IN timings change from first IN to last IN; i.e. we expected weak or no significant correlations between the first IN and song and strong correlations between the last IN and song. Additionally, we expected this pattern to be present in greater than 50% of the birds. Contrary to this expectation, we found significant correlations in timing of last IN and timing of song in only 6/20 birds and these correlations were weak (median - 0.14; range: 0.06 - 0.2; Table 1). Only 1/20 birds had significant correlations between first IN timing and song timing. Overall, a majority of birds did not show significant correlations between IN and song intervals or improvement in correlations with song interval from first to last IN intervals suggesting that changes in IN timing do not represent motor preparation for song syllable timing.

### IN acoustic features are correlated with song features

Similar to timing, we examined correlations between the acoustic features of INs and the acoustic features of the first song syllable (see Fig. 3 for an example). Log amplitude and mean frequency of the last IN were correlated with the corresponding features of the first song syllable in a large proportion of birds (Fig. 4, Table 1, p < 0.05, Pearson’s correlation co-efficient). Surprisingly, we also found significant correlations between the acoustic features of the first IN and the first song syllable in most of these birds (Fig. 4, Table 1, p < 0.05, Pearson’s correlation coefficient), albeit the strength of the correlations were slightly lower when compared to the strength of correlations between the last IN and the first song syllable (Fig. 4, p = 0.004, Mean Frequency and p=0.015, Log Amplitude for comparisons of r-values). The presence of correlations with the first IN (Fig. 4, Table 1) suggested the possibility that these correlations could alternatively be related to previously described global correlations between syllables within individual bouts (Glaze and Troyer, 2006). In support of this idea, we also found significant correlations between the acoustic features of the first two song syllables (Fig. 4, Table 1, p < 0.05, Pearson’s correlation co-efficient). The correlations were similar in strength to the correlations between IN features and song features and were present in a similar proportion of birds (Table 1, Fig. 4, p > 0.05, Wilcoxon signed-rank test). In a majority of birds, significant correlations were also absent for other acoustic features related to syllable identity (duration, pitch goodness, frequency modulation), further supporting the idea that syllable feature correlations represent global control of IN and song syllables, rather than INs reflecting motor preparation for the upcoming song.

To rule out the possibility that such correlations arose because the distance from the suspended microphone to the bird varied from bout-to-bout, we also examined similar correlations in a dataset from a different set of birds with head-implanted microphones (Suri and Rajan, 2018). In these birds, the distance between the microphone and the bird was always fixed. We found similar correlations in a similar proportion of head-implanted birds (Table 1) confirming that these correlations were not just due to differences in relative distance from the microphone.

To better understand the origin of these correlations of globally controlled features, we further analysed song recordings from birds with bilateral cuts in the tracheosyringeal nerve, the nerve that carries neural input to the syringeal muscles. In these birds, syllable identities are lost as syllables are reduced to harmonic stacks, but the temporal patterning of song remains similar to pre-nerve cut as this is controlled by respiratory motor neurons (Bottjer and Arnold, 1984; Vicario, 1991; Williams and McKibben, 1992; Roy and Mooney, 2007). Significant correlations in most acoustic features were present in a similar proportion of these birds (Table 1, p < 0.05, Pearson’s correlation co-efficient). These results suggest that bout-to-bout correlations in acoustic features of INs and song syllables are driven by bout-to-bout differences in the control of respiratory pressure.

Overall, these results show the presence of significant correlations in IN acoustic features (first and last IN) with acoustic features of the first song syllable. Additionally, correlations between acoustic features of the first two song syllables suggest that the correlations between IN features and first song syllable features are not a result of motor preparatory function of INs but, instead, reflect the fact that INs are also vocalizations that are part of the song sequence and are controlled by the same neural pathways.

### Song and INs show age-related changes

We next examined a second prediction of the motor preparation hypothesis, age-related song changes and their correlation with IN properties. Previous studies have shown age-related changes in song tempo and song stereotypy; songs become faster and more stereotyped as birds get older (Brainard and Doupe, 2001; Pytte et al., 2007; Glaze and Troyer, 2013; James and Sakata, 2019). If INs represent motor preparation, we would predict simultaneous changes in specific IN properties with age. In addition, we would expect age-related changes to be predicted by correlations between these IN properties and song properties at an earlier age. For instance, if IN number increases with age and songs get more stereotyped with age, we would expect higher IN number to be associated with more stereotyped songs, on a bout-to-bout basis, at an earlier age (Fig. 1D).

To test this, we first recorded birds at multiple time-points (median - 3; range - 2-8 time-points per bird) from ∼90 days post-hatch to ∼3 years of age (Fig. 5A). As compared to the first day of recordings, we found increases in IN number in most birds (Fig. 5A, red circles represent birds with increases over age, black circles represent birds with decreases over age and gray circles represent birds that do not change over age). We divided our recordings into 3 age categories, namely, (1) within 1 year post hatch, (2) 1-2 years post hatch and, (3) > 2 years post hatch. For some of the birds we had multiple recordings within each of these categories, typically within a few days of each other. For each bird we calculated pair-wise differences in mean IN number between the different age categories. Differences in mean IN number were largest when the first recording day was within 1 year post hatch and the second recording day greater than 1 year (Fig. 5B, bigger circles for Age 1 < 365 dph and Age 2 > 365 dph). For each bird, we chose pairs of days between age categories with the largest difference in mean IN number (see Methods for details, Fig. 5B, ’*’ represents days chosen for further analysis) and then used changes between these pairs of days to further compare age-related changes across birds. We chose this strategy to increase our chances of testing the prediction of the motor preparation hypothesis as we expected greater correlations for these days as mentioned above. All of the analyses described below were carried out with these days.

Mean IN number and the associated “time-to-song” increased in the 1st year post-hatch (Fig. 6C, p = 0.02, Kruskal-Wallis test, p = 0.09 for <1 yr to > 1yr vs. day-day and p = 0.02 for <1 yr to > 1 yr vs. > 1yr, post-hoc Tukey-Kramer test, Fig. 6D, p=0.003 Kruskal-Wallis test, p=0.04 for day-day and <YR to >1yr, p=0.003 for <1YR to >yr and >1yr, post-hoc Tukey-Kramer Test). Song tempo, for the first song motif (that followed the INs at the beginning) in the bout, increased significantly, as seen by the significant shortening of first song motif duration (Fig. 6B, p = 0.007, Kruskal-Wallis test, p = 0.005, day-day vs. <1YR to > 1yr, post-hoc Tukey-Kramer test). In our birds, song stereotypy did not increase significantly across the 3 age-groups, although song structure itself significantly changed in the first year (Table 2, song motif similarity index and motif temporal similarity between days). Other features of songs that showed changes in the first year were reduced sequence entropy, increased pitch goodness and reduced variability in frequency modulation of first song syllable. Significant changes were also observed for acoustic features of INs with increased frequency modulation, decreased CV of duration, pitch goodness and increased CV of frequency modulation of INs. We also observed some trends that approached significance including longer intervals between the first two INs, increased pitch goodness of INs, increased number of motifs in a bout, and reduced duration of the first motif syllable (Table 2, p>0.05 and <0.1). Overall, these results document age-related changes in IN properties that occur along with age-related song changes.

**Table 2:**
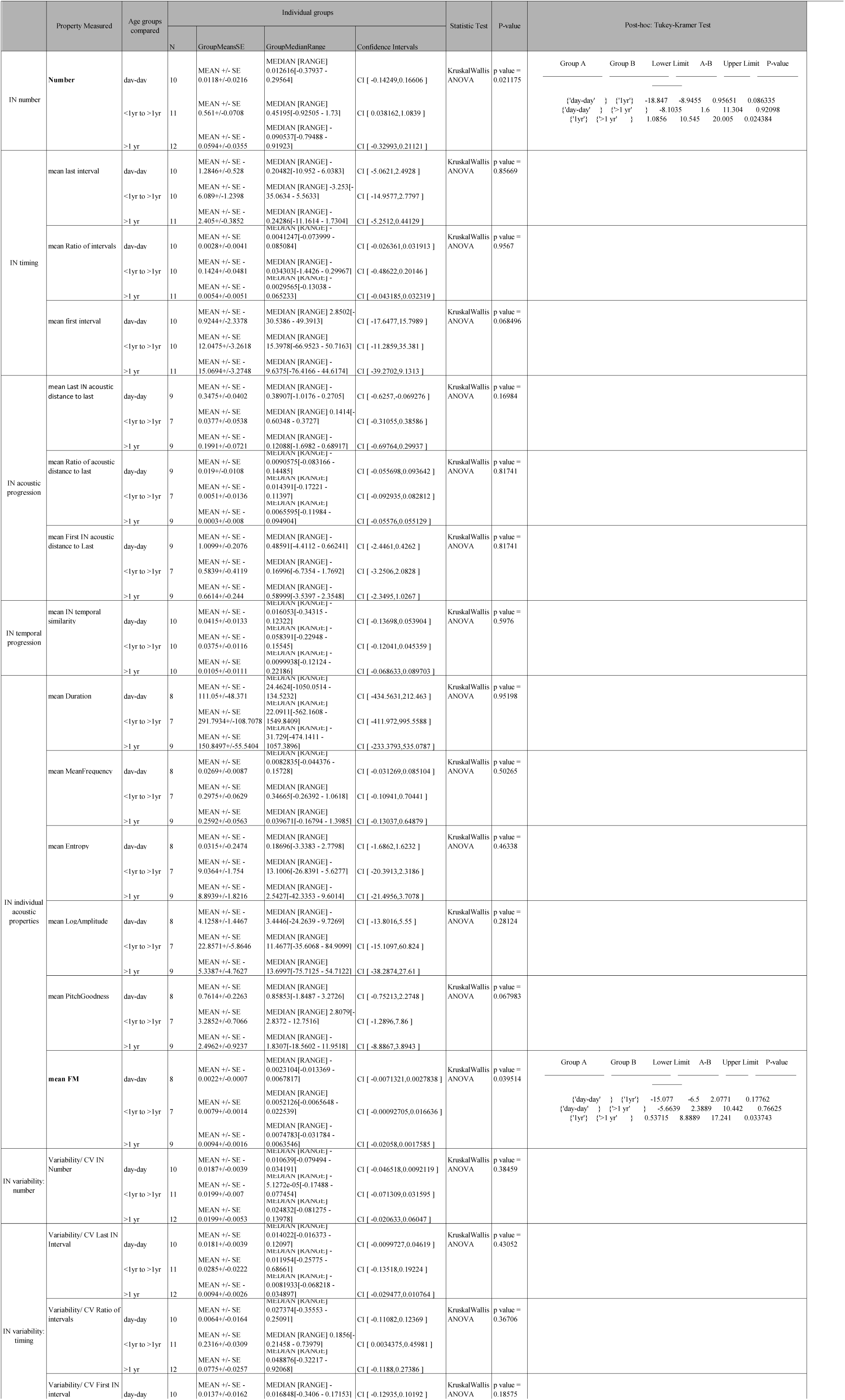
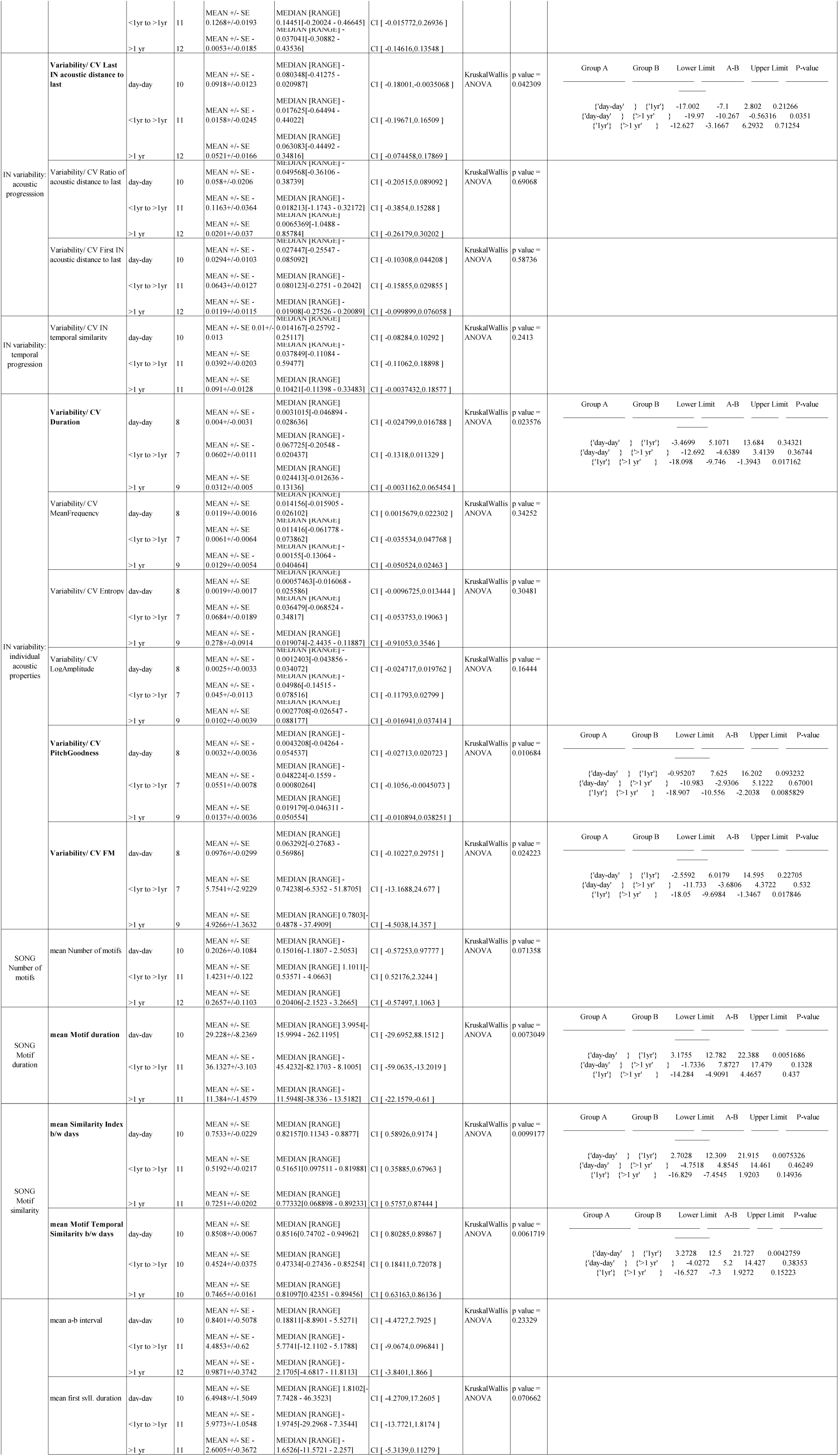
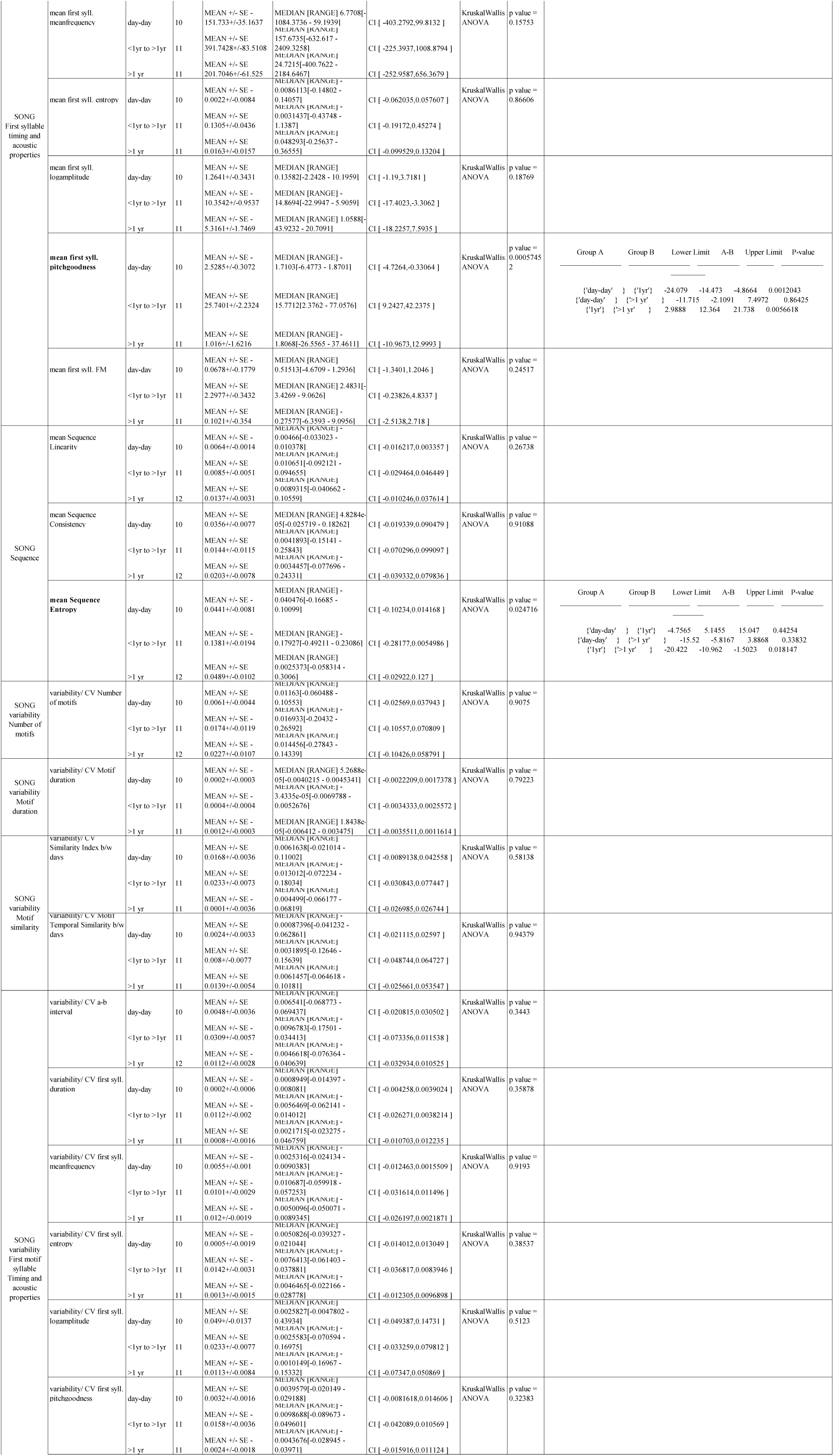
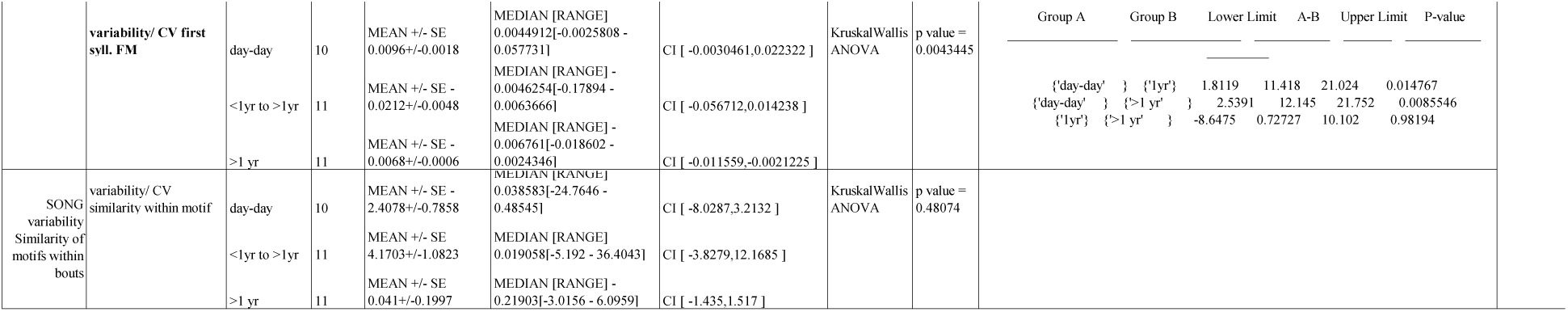
IN and Song changes with age. Changes in mean or variability of IN properties between pairs of days belonging to 3 age categories: day-day, <1YR to >1yr and > 1 yr. All pairs of days are the same pairs of days selected for maximum change in IN number in Fig. 5. Names of properties that significantly change with age are highlighted in bold.

### IN features that change with age do not show bout-to-bout correlations with song features that change with age

If INs represented motor preparation, the correlations between IN features and song features on a bout-to-bout basis should predict long-term changes in both IN and song features (Fig. 1D). For instance, our results demonstrated an age-related speeding up of song (decrease in song motif duration) and an age-related increase in mean IN number with age. If INs were preparatory, then these changes would be predicted by an already existing negative correlation between the number of INs in a bout and song motif duration. However, we did not find any correlation between the number of INs in a bout or the time-to-song and song motif duration (Fig. 7A, 7B, p = 0.86, Kruskal-Wallis Test and p = 0.88, Wilcoxon signed-rank test respectively). We also compared first song motif stereotypy between bouts with fewer INs than the median IN number and bouts with IN number greater than the median IN number. First song motif stereotypy was not significantly different (Fig. 7C, 7D, p=0.44.

**Fig. 7.**
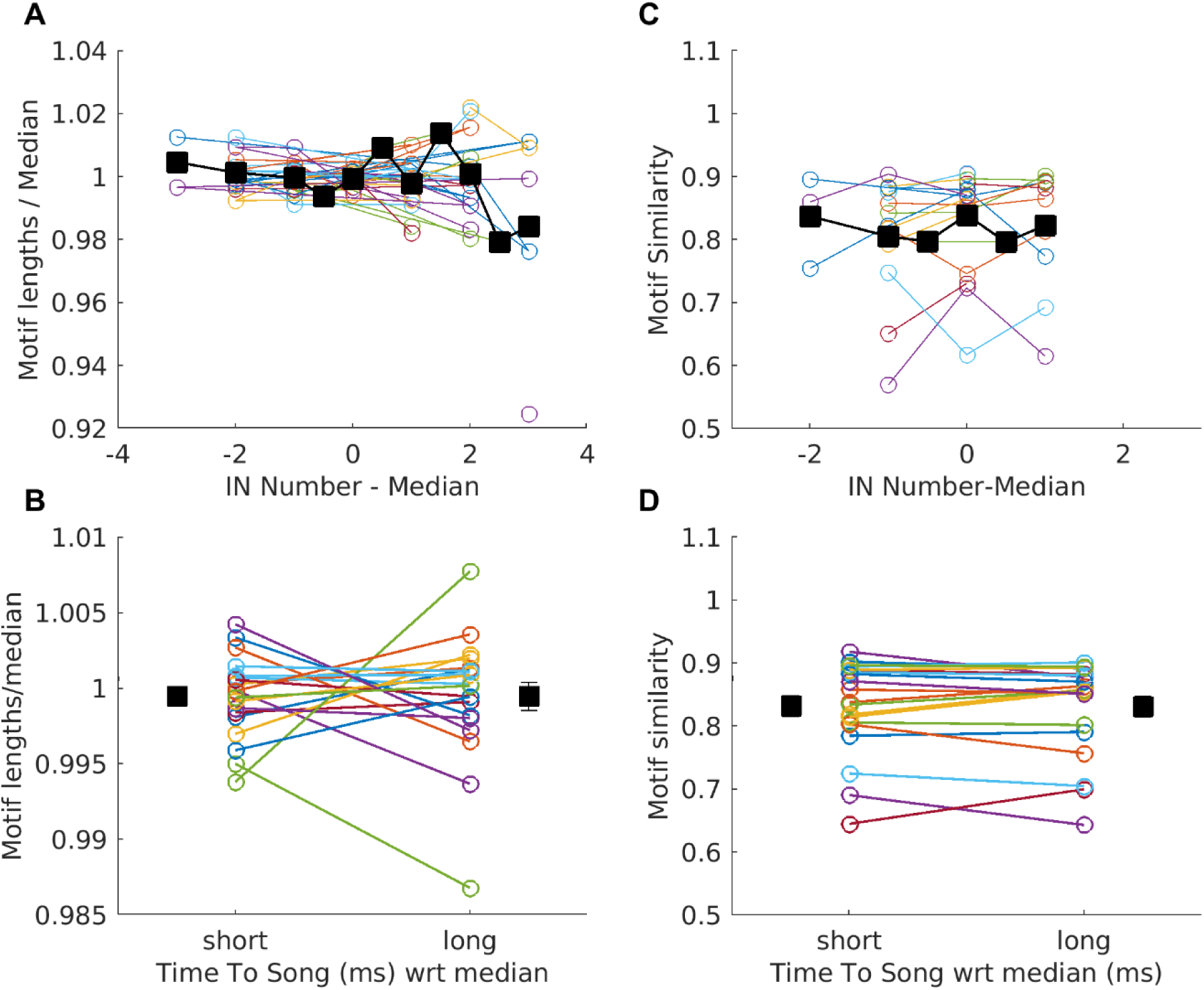
Correlated IN and song changes with age are not related within a session. (A, C) Motif length normalized to median length (A) and motif similarity (C) are plotted for different IN numbers relative to the median IN number. Black squares represent mean across birds. (B, D) Motif length normalized to median length (A) and motif similarity (C) are plotted for short and long time-to-song. Squares and whiskers represent the mean and SEM across birds. In all plots, circles joined by lines represent mean values for one session from individual birds and different colours represent different birds. p > 0.05 KruskalWallis ANOVA in (A) and (C), p > 0.05, Wilcoxon signed-rank test in (B) and (D).

Kruskal-Wallis Test and p=0.65, Wilcoxon signed-rank test respectively).

Similarly, we did not find any bout-to-bout correlations between other IN features that change with age and song features that change with age (Table 3). Overall, these results demonstrate that song features and IN features change independently with age. The absence of correlations between features that change do not satisfy the predictions of the motor preparation hypothesis. Rather, these changes demonstrate independent age-related changes in both INs and songs, suggesting the possibility that age affects neural circuits controlling INs and songs.

**Table 3.**
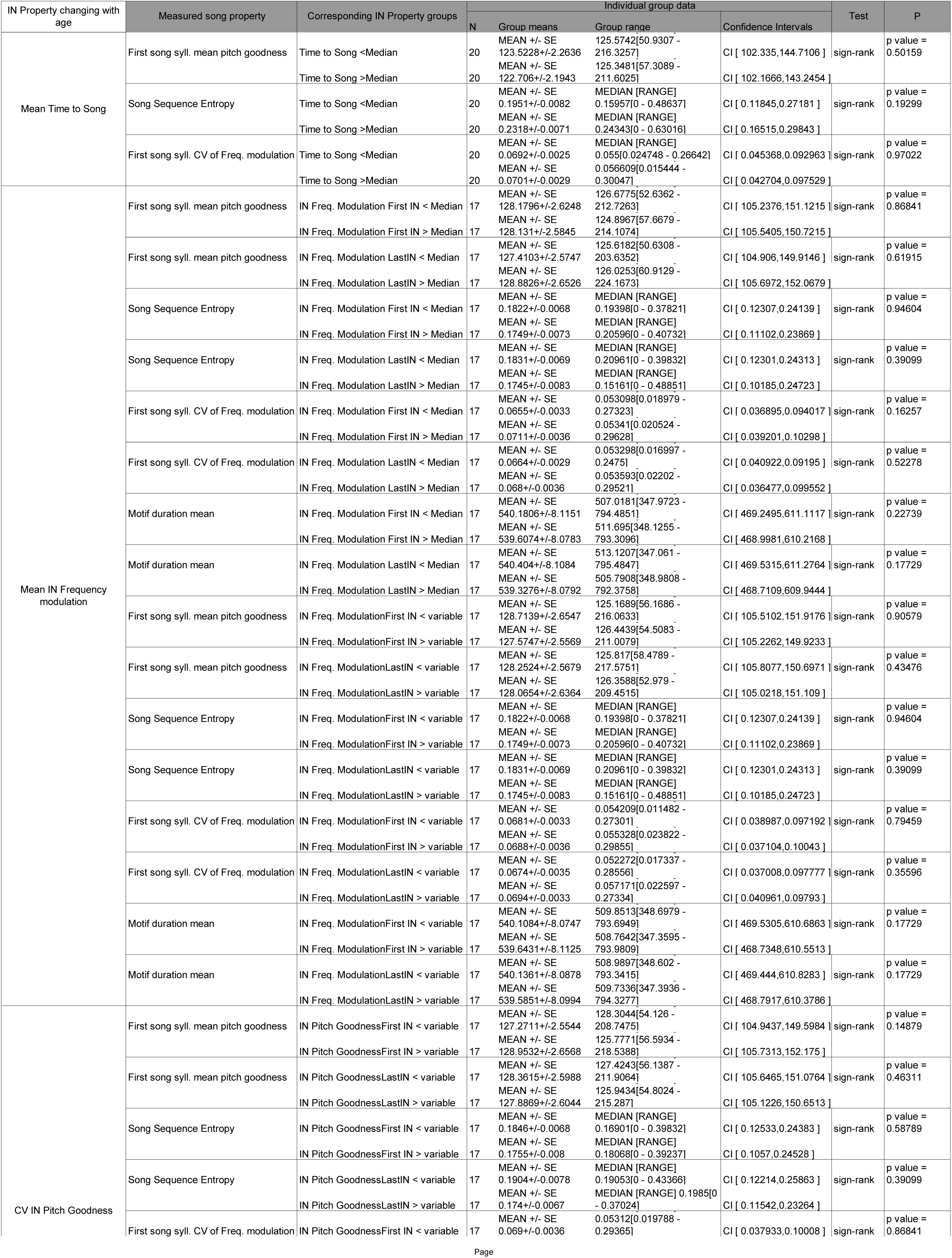
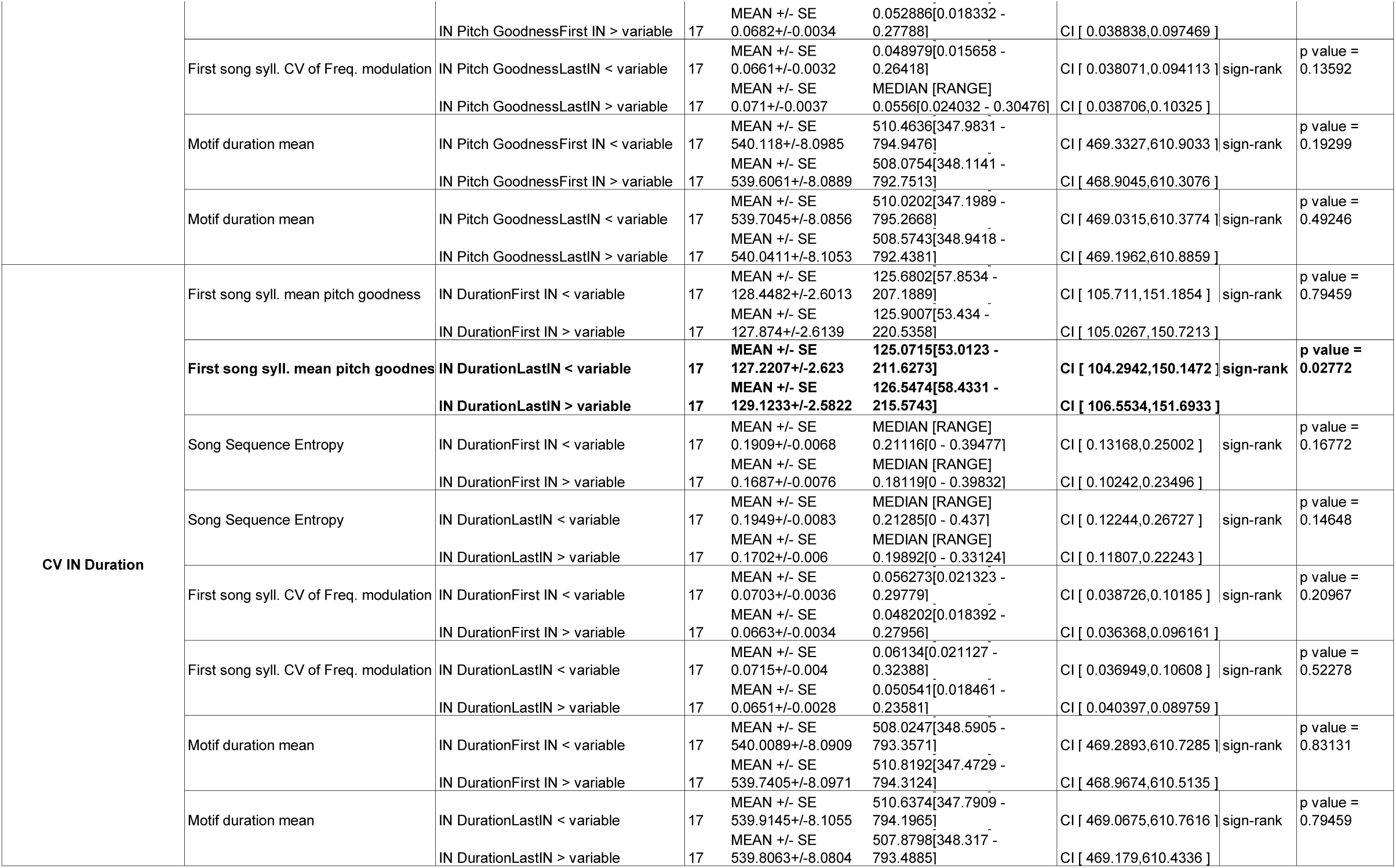
Within-day relationship of IN and Song properties that change with age. For IN properties (Time to song, frequency modulation, CV of pitch goodness and duration) and song properties (Motif duration, mean pitch goodness of first song syllable, sequence entropy of the motif, CV of frequency modulation of first song syllable) that change with age, changes within-day were compared. The bouts were split based on mean or CV of IN property into two groups and corresponding mean or CV of song property was measured within each bird. The paired-groups were tested for statistical differences using Wilcoxon sign-rank test.

### IN acoustic features share similarities with other syllable repeats but speeding up of intervals between INs is unique to IN repeats

Our current results suggest that none of the IN properties, namely, number, timing and, acoustic features represent motor preparation for features of the upcoming first song syllable. Further, IN acoustic features co-vary with song syllable acoustic features. One possibility for this correlation in acoustic features is the fact that INs are also vocalizations like song syllables. In fact, a recent study has shown that mean IN number and IN acoustic features are learned by young zebra finches, similar to learning of song syllables (Kalra et al., 2021).

Unlike most song syllables, INs repeat and this repetition may drive some of the changes in properties of INs as they progress to song. To examine this possibility, we next asked if the properties of INs are more similar to the properties of song syllables that repeat. Such song syllable repeats are present in a small fraction of zebra finches (Fig. 8A top; example spectrogram with repeat of the song syllable ’c’). Zebra finches also repeat calls (other non-song vocalizations that can be produced outside of song bouts for communication) outside of their song bouts (Fig. 8A bottom). To compare repeats across these three categories, we chose birds with motif syllable repeats. Both mean IN number and the variability of IN number from bout to bout were not significantly different from the mean number and variability of repeats for other categories (Fig. 8B, p=0.049 Kruskal-Wallis test followed by post-hoc Tukey-Kramer test, p = 0.31 for IN vs. song and p =0.67 for IN vs. calls, Fig. 8C p=0.20, Kruskal-Wallis Test).

**Fig. 8.**
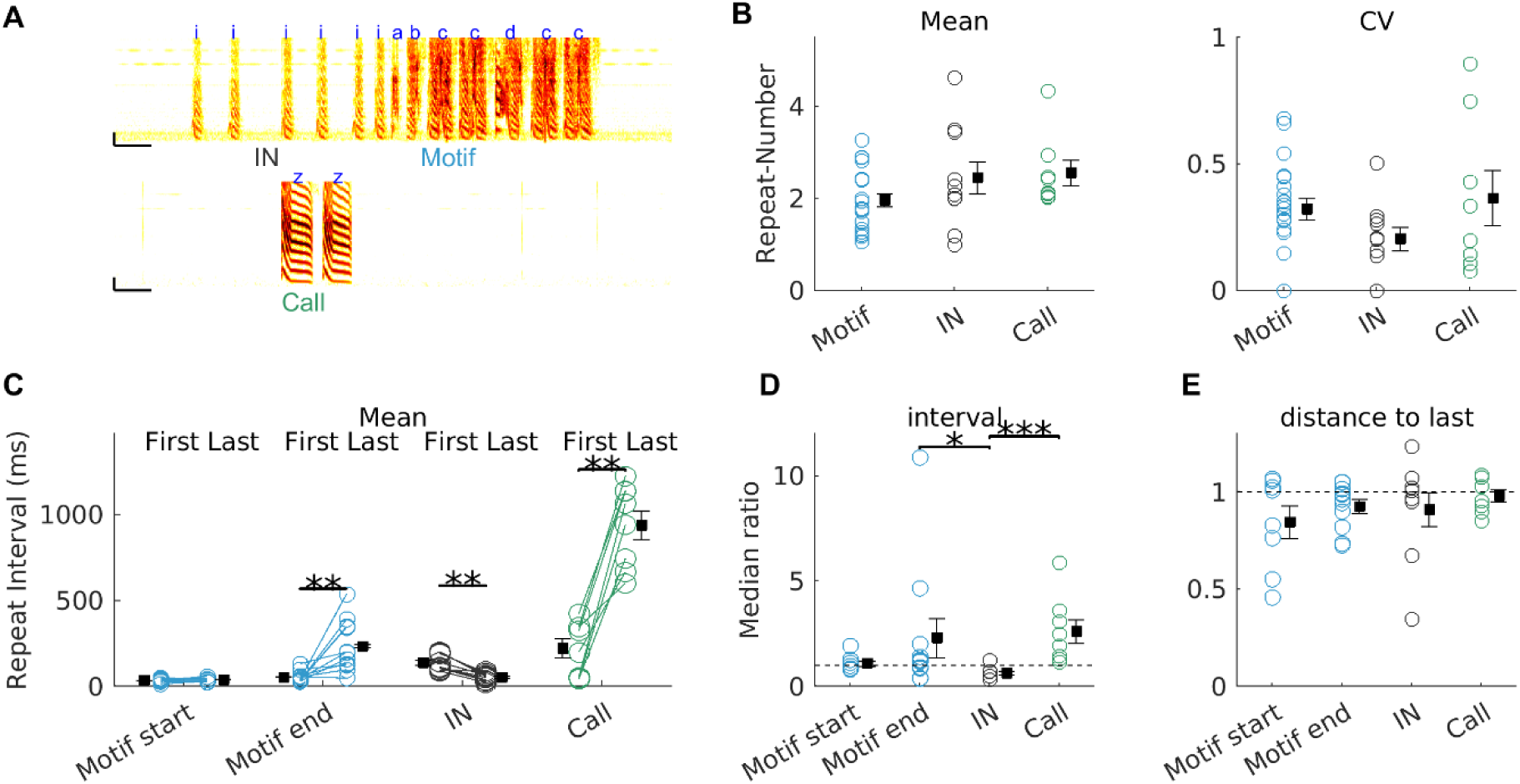
Temporal progression of INs is not present in other repeats. (A) Top - example spectrogram of a bird that produces repeating INs ‘i’ and repeating syllable ‘c’ within the motif. Bottom – example of a bird producing repeats of call syllables ‘z’ outside song. (B-E) Comparison of repeat number (B left), repeat number variability (B right), repeat interval (C), median ratio of repeat interval (D), and median ratio of acoustic features of successive syllables (E) between different types of repeats (motif repeats, IN repeats or call repeats). Circles represent data from individual birds, squares and whiskers represent the mean and SEM across birds. In C, lines join data from the same repeat syllable across different positions. ** denotes *p* < 0.01 Wilcoxon signed-rank test in (C), * denotes *p* < 0.05, *** p denotes < 0.005, Kruskal-Wallis test, followed by posthoc Tukey-Kramer Test in (D). Dashed lines in D and E represent a ratio of 1.

Interestingly, intervals between repeat syllables showed interesting differences between the three categories of repeats. For all categories of repeats, we compared the mean interval between the first two syllables in the repeat sequence (first interval) and the last syllable of the repeat sequence and the next syllable in the bout (last interval). As shown earlier (Rajan and Doupe, 2013; Rao et al., 2019), for IN repeats, the last interval was significantly shorter than the first interval (p=0.004, Wilcoxon signed-rank test) and the opposite was true for call repeats with the last interval being significantly longer than the first interval (Fig. 8C, p=0.008, Wilcoxon signed-rank test). Interestingly, for motif syllable repeats, we observed differences depending on the position of the repeat within the song motif; for repeats that were the last syllable of the song motif, the last interval was significantly longer than the first interval (p=0.002, Wilcoxon signed-rank test), while for repeats that occurred at the beginning of the song motif, the last and first interval were not significantly different (Fig. 8C). Thus, the speeding up of intervals between successive INs was a unique feature of INs (Fig. 8D - ratio of successive intervals is less than 1 only for IN repeats; mean +/- SE - 0.65+/-0.03). The acoustic features of all repeats changed from first to last syllable of the repeat and was not significantly different for any of the categories (Fig. 8E, p=0.71), suggesting that changes in acoustic features were a property of syllable repetition. Overall, these results show that bout-to-bout variation in number of INs and variation in acoustic properties of successive INs are a property of all syllable repeats. However, the speeding up of intervals between successive INs is not a property of all syllable repeats but is unique to INs at the beginning of the bout.

## Discussion

Here, we tested two predictions of the hypothesis that the variable repeats of INs that precede song represent motor preparation for the upcoming song. First, we examined bout-to-bout correlations between the first song syllable and three properties of INs, namely, (1) number before song, (2) timing of INs and (3) acoustic features of INs. Both number and timing of INs were not correlated with features of the first song syllable. Log amplitude and mean frequency of the last IN were correlated with the corresponding acoustic features of the first song syllable. However, acoustic features of the first song syllable were also correlated with those of the first IN and the second song syllable. This suggested that INs are also vocalizations that are part of the song sequence and IN-song correlations reflect global bout-to-bout variation in acoustic features. Second, we found an age-related increase in mean IN number along with an age-related increase in the speed of song. Age-related changes in INs and song were independent and were not predicted by pre-existing bout-to-bout IN-song correlations in the same sets of features. Finally, we compared IN repeats with other types of syllable repeats. Just like IN repeats, other syllable repeats also varied in number between bouts and varied in acoustic properties as the repeats progressed. The speeding up of intervals between INs with each successive repeat was unique to IN repeats, while the intervals between other kinds of syllable repeats remained constant or increased with each successive repeat. Overall, these results show that INs and song syllables share similarities in acoustic features that suggest shared neural control, while IN and song syllable timing are different highlighting possible differences in the neural control of timing.

### Song control pathways may control changes in IN acoustic features

Our results show correlations between IN acoustic properties and the corresponding acoustic properties of the first song syllable. While, this is expected from the motor preparation hypothesis, we also found correlations between the first IN and the first song syllable and the first and second song syllables, suggesting the presence of global correlations between syllables within a bout. Similar correlations in the duration of syllables within a song motif have been described earlier (Glaze and Troyer, 2006). Our results show that such correlations extend to INs as well, suggesting that INs are also syllables similar to song syllables.

Consistent with this idea, we also found changes in acoustic features for song syllables that repeat within song and non-song vocalizations that repeat outside of song sequences. These results suggest that INs and song syllables are controlled by the same neural circuitry that control song syllable production.

What areas of the brain control song syllable and IN production? Song syllable structure and timing are known to be controlled by the song motor pathway consisting of premotor nucleus HVC (used as a proper name) and its projection to the robust nucleus of the arcopallium (RA) (Hahnloser et al., 2002; Long and Fee, 2008; Fee and Scharff, 2010). There is some evidence that INs might also be controlled by the same circuitry, albeit with some differences. HVC neurons and RA neurons show temporally precise bursts of neural activity during song syllables (McCasland, 1987; Yu and Margoliash, 1996; Hahnloser et al., 2002; Kozhevnikov and Fee, 2007; Long et al., 2010) and some of these neurons also show precise bursts of neural activity during INs (Hahnloser et al., 2002; Kozhevnikov and Fee, 2007; Rajan and Doupe, 2013). However, some HVC neurons also show differences in activity depending on the position of INs within the sequence (Rajan and Doupe, 2013) similar to HVC firing in other songbird species with more complex songs with syllable repeats (Fujimoto et al., 2011; Cohen et al., 2020). Cooling of HVC slows down song syllables at multiple timescales and cooling of HVC also slows down INs, albeit by 70% of the extent of slowing down of song syllables (Long and Fee, 2008). Finally, complete bilateral lesions of HVC abolish the normal production of song syllables (Nottebohm et al., 1976; Simpson and Vicario, 1990; Aronov et al., 2008; Chen et al., 2014), but whether they entirely abolish INs is unclear. Outside of the song motor pathway, a recent study has also shown that lesions of the midbrain dopaminergic nucleus A11 makes birds completely mute for song (Ben-Tov et al., 2023). Given the soft amplitude of initial INs, it is unclear whether these birds produce a few INs that do not progress normally to song. Overall, our results and the results of earlier studies suggest that IN production is also partly controlled by the song motor pathway and partly controlled by neural circuits outside of the song motor pathway.

### What controls the speeding up of gaps between INs?

Our results show that IN timing was not correlated with the timing of the upcoming song syllables. However, IN timing, characterized by a speeding up of intervals between successive INs, was not a feature of all other syllables that repeat. Infact, for other syllables that repeated, intervals between successive syllables remained the same or became longer. This suggests that the speeding up of intervals between INs could indicate the time to song initiation (Rao et al., 2019). In primates, one aspect of neural preparatory activity is the consistent, and strong, correlation with the time at which the movement is initiated (Kaufman et al., 2016). Similarly, the speeding up of intervals between INs could reflect the readiness of specific brain regions to initiate song.

Which brain regions could be involved in this process? One candidate region is the premotor nucleus HVC and its inputs. As mentioned above, HVC controls the temporal dynamics of syllable production and is necessary for normal song production (Nottebohm et al., 1976; Simpson and Vicario, 1990; Aronov et al., 2008; Long and Fee, 2008). Intracellular recordings have shown that HVC_RA_ neurons (HVC neurons projecting to motor nucleus RA) are more depolarised during singing compared to non-singing and this depolarisation begins during the INs (Long et al., 2010). This depolarisation could potentially come from dopaminergic input from midbrain dopaminergic nucleus, A11. In support of the idea that this dopaminergic input is important for IN progression to song, dopaminergic blockers in HVC make birds sing strings of INs, without progressing to song, directed towards female birds (Ben-Tov et al., 2023).

How could this depolarisation of HVC neurons during INs drive IN progression to song? Initiation of song syllables is believed to be driven by thalamic input, from nucleus Uvaeformis (UVa), to HVC at the start of each song syllable (Moll et al., 2023). UVa neurons are also active at the onset of INs suggesting the possibility that UVa input drives HVC at the start of each IN (Danish et al., 2017). The depolarisation state of HVC could play a role in determining the time when UVa input successfully initiates an IN; more depolarised state in HVC, sooner HVC neurons are activated and this in-turn could lead to a shorter interval between INs. In this model, the speeding up of INs would reflect the gradual reduction in the time taken by HVC neurons to respond to UVa input. This model also predicts that Ins would not be necessary when HVC neurons are already depolarised. This is supported by the fact that successive songs within a bout can be initiated without INs (Sossinka and Böhner, 1980; Rajan and Doupe, 2013).

### Do INs have a preparatory function?

Our results do not completely support the predictions of the motor preparation hypothesis postulated based on neural activity in primates. This suggests that the parallels between INs reaching a consistent “state” before song initiation and primate preparatory neural activity reaching a consistent neural state before movement initiation are restricted to their correlations with the time to movement initiation. This hypothesis predicts that disruption of INs and IN-related activity in HVC would only delay song initiation without affecting the features of upcoming song. Further studies disrupting IN-related neural activity in HVC could be used to test this.

Second, our results showing the similarities in the acoustic features of INs and other repeated syllables suggest that INs might be a special form of repeating syllables that occur at the beginning of zebra finch song bouts. Other songbird species with more complex song bouts have syllable repeats that occur regularly within the song (Okanoya, 2004; Yarden et al., n.d.). However, two questions remain unanswered, namely, (1) do IN-like repeating syllables occur at the beginning of song bouts of other songbird species and (2) do the properties of IN repeats share similarities with syllables repeats that are present in the song sequences of other songbird species. Further comparative analysis of song bouts across different songbird species could be used to address these questions.

Overall, our results suggest that IN acoustic structure and repetition share similarities with song syllables, suggesting shared neural control of IN acoustic structure and repeat number. The speeding up of intervals between INs is a unique feature of INs and could reflect different neural mechanisms controlling the timing of INs.

## Acknowledgments

We would like to thank Prakash Raut for bird care. We thank Shikha Kalra, Ananya Kumar, Aditi Agarwal, Vishruta Yawatkar, Harsha K Kumar, Sharvari Tamhankar, Gaurav Isola for labeled songs or songs recorded at different ages, and Harini Suri for sharing head-implanted microphone recordings. We thank Allison Doupe and Michael Brainard for song recordings of birds with repeat syllables. We also thank Mimi Kao, Michael Long and members of the Rajan lab for useful discussions. DR is currently at Department of Neuroscience, University of Copenhagen, Blegdamsvej 3B, Denmark 2200, Copenhagen N

**Extended Data Table supporting Figure 2**

Statistical test results for data from Fig 2. - Fig. 8

